# Local adaptation of *Aedes aegypti* mosquitoes to *Wolbachia*-induced fitness costs

**DOI:** 10.1101/2022.05.06.490959

**Authors:** Perran A. Ross, Ary A. Hoffmann

## Abstract

*Aedes aegypti* mosquito eggs can remain quiescent for many months before hatching, allowing populations to persist through unfavorable conditions. *Aedes aegypti* infected with the *Wolbachia* strain *w*Mel have been released in tropical and subtropical regions for dengue control. *w*Mel reduces the viability of quiescent eggs, but this physiological cost might be expected to evolve in natural mosquito populations that frequently experience stressful conditions. We therefore compared the costs of *w*Mel infection for quiescent egg viability in field-derived and laboratory populations. Quiescent egg viability was highly variable in *w*Mel-infected populations, with greater costs of *w*Mel in field-derived populations. In contrast, there was little variation between matched field-derived and long-term laboratory populations lacking *w*Mel, suggesting that laboratory adaptation does not influence this trait and that differences are due to *w*Mel infection. Comparisons of populations collected a year apart show a decline in costs under laboratory rearing conditions involving a rapid turnover of mosquito generations; this pattern was consistent across populations despite their origin, suggesting adaptation of mosquitoes to the *w*Mel infection under laboratory conditions. Reciprocal crossing experiments confirm that differences in quiescent egg viability were mainly due to the genetic background and not *Wolbachia* alone. *w*Mel-infected mosquitoes hatching from long-term quiescent eggs showed partial loss of cytoplasmic incompatibility and female infertility, highlighting additional costs of long-term quiescence. Our study provides the first evidence for a shift in *Wolbachia* phenotypic effects following deliberate field release and establishment and it highlights interactions between *Wolbachia* infections and local adaptation. The unexpected changes in fitness costs observed here suggest potential tradeoffs with undescribed fitness benefits of the *w*Mel infection.

## Introduction

*Aedes aegypti* is well adapted to anthropogenic environments across the world, that include not only the ability to tolerate pesticides in these environments [1] but also life history strategies that allow it to breed under variable conditions [2]. One adaptation to variable conditions is the ability for its eggs to enter a quiescent stage, where they remain viable for periods without water before hatching following rainfall or human-associated activities that increase water availability [3]. This phenotype is likely to be particularly important in climates with an extended dry season where populations may otherwise not persist [4]. *Aedes aegypti* show local adaptation to quiescence [5], where populations from locations that experience longer dry seasons or infrequent water availability have longer viability as quiescence eggs.

While egg quiescence is expected to contribute to fitness under many natural conditions, it will be less important in the laboratory, at least when mosquitoes are maintained under conditions that lead to a constant turnover of generations that maximize population productivity. This raises the issue of whether mosquito populations under constant turnover might adapt to laboratory conditions in terms of quiescence as well as other traits like mating behavior and stress tolerance [6]. While there will often be a lack of selection to maintain long-term quiescent egg viability, effects of laboratory adaptation on this phenotype have rarely been tested [5].

*Wolbachia* endosymbionts are being released in various locations around the world and having substantial localized impacts on dengue transmission [7-9]. One challenge is that *Wolbachia* can have a substantial impact on quiescent egg viability; this was first recognized for the *w*MelPop *Wolbachia* infection [10, 11] but also applies to *w*AlbB [12, 13]. These costs may have contributed to unsuccessful establishment of both *w*MelPop and *w*AlbB in some locations [8, 14]. In fact, quiescent egg viability costs have been proposed as a tool for seasonal population suppression [4, 15]. The widely released *w*Mel strain also reduces egg viability [16, 17] but the effects are usually weaker compared to other strains, and this has not prevented its establishment in several locations, even in places with long dry seasons such as Cairns, Australia [18].

Releasing a *Wolbachia* strain that induces fitness costs is expected to cause evolutionary changes in either the mosquito or *Wolbachia* to attenuate these effects [19, 20]. While the *w*Mel genome has remained unchanged [21-23] there is currently a limited understanding mosquito adaptation to *Wolbachia* infections. The *Ae. aegypti* genome shows minor changes across the decade in which *Wolbachia* have been released in North Queensland, Australia and some of these changes might relate to fitness effects [24]. Selection experiments show that the quiescent egg viability costs of *w*MelPop can be ameliorated through selection due to changes in the nuclear genome [15] and costs to some traits also appear to have shifted over time though long-term lab rearing [25].

Currently, there is no evidence that *w*Mel fitness costs have shifted much under field conditions [23, 26]. There is also no evidence for differences in fitness costs of *w*Mel between mosquito genetic backgrounds that have had long-term coadaptation versus naïve backgrounds [23]. However, phenotypic comparisons have covered only a limited number of traits, most of which are only minimally influenced by *w*Mel infection, and quiescent egg viability costs have not been compared between backgrounds. In field populations, we might expect to see *Wolbachia* costs associated with quiescence become weaker over time. This could have impacts on population dynamics; *w*Mel is expected to reduce the *Ae. aegypti* population size relative to competing container species due to its fitness costs [27], thus indirectly reducing virus transmission, but evolution could mitigate this benefit. Data from long-term post-release monitoring of *w*Mel in Yogyakarta, Indonesia suggest minor or absent fitness costs of *w*Mel infection under these conditions, as the ratio of *Ae. aegypti* to competing *Ae. albopictus* decreased only marginally [28].

In this study, we measured the effects of *w*Mel infection on *Ae. aegypti* quiescent egg viability in mosquito populations originating from different locations. We then performed reciprocal backcrossing to identify the cause of differences in fitness costs. Costs were driven strongly by mosquito genetic backgrounds, with surprisingly higher costs in field-collected mosquitoes. In the absence of *Wolbachia*, there was no evidence for evolution of quiescent egg viability in long-term laboratory populations despite the lack of selection to maintain this trait. The differential fitness costs of *w*Mel in the backgrounds observed here provides some of the first direct evidence of mosquito adaptation to novel *Wolbachia* infections. We also identify additional cross-generational costs of *w*Mel in quiescent eggs. Our results have implications for the spread of *Wolbachia* in different local environments, and the use of laboratory studies to test fitness costs of *Wolbachia*.

## Methods

### Ethics statement

Blood feeding of female mosquitoes on human volunteers for this research was approved by the University of Melbourne Human Ethics Committee (approval 0723847). All adult subjects provided informed written consent (no children were involved).

### Field collections and colony maintenance

We established *Ae. aegypti* populations from eggs collected from ovitraps placed in suburban Cairns in 2016, 2018, 2019 and 2020 as described previously [29]. Collections focused on two suburbs: Yorkeys Knob (YK) and Gordonvale (GV), where *w*Mel-infected *Ae. aegypti* were first released in 2011 [18], as well as suburbs in central Cairns, where staggered releases took place from 2014 [7]. Each *Ae. aegypti* population was established from a pool of 15-50 ovitraps and at least 200 founding individuals. Laboratory populations were maintained at a census size of ∼450 individuals each generation at 26°C and a 12:12 light:dark cycle as described previously [30]. Female mosquitoes were fed on the forearm of a single human volunteer for egg production. Repeated collections from the field enabled us to establish *w*Mel-infected populations from the same origin but with different numbers of generations of laboratory rearing, with approximately 12 generations per year. We also performed experiments on the same populations at different times allowing us to track changes over time, though experiments were not directly comparable due to potential differences in egg storage conditions. In all experiments, we included a long-term laboratory population (*w*Mel Lab), which was collected from Cairns in 2014 and had spent at least 60 generations in the lab at the time of the first experiment.

### Antibiotic curing

To generate *w*Mel-infected and uninfected mosquitoes with matching genetic backgrounds, populations were split in two, with one population treated with tetracycline and the other left untreated. Treated populations were provided with 10% sucrose containing 2 mg/ mL of tetracycline hydrochloride for two consecutive generations, followed by two generations of recovery (with no tetracycline) before experiments. Populations collected from the field began treatment within the first 1-2 generations in the laboratory to minimize potential effects of laboratory adaptation. In the generation following treatment, 30 individuals from each treated population and untreated population were screened for the presence of *w*Mel according to previously described methods [31]. Populations were only used in experiments if *w*Mel was absent from all individuals in treated populations and present in all individuals in untreated populations. *w*Mel-infected populations were cured each time they were used in experiments; *w*Mel-infected and uninfected counterparts were not maintained as separate populations for more than 5 generations to minimize potential effects of genetic drift [32].

### Quiescent egg viability

We measured the quiescent egg viability of *Ae. aegypti* in several experiments using a standardized approach. Females in colony cages (aged 5-7d, starved of sugar for 1 d) were blood fed and six plastic cups (250 mL) filled with larval rearing water and lined with sandpaper strips (Norton Master Painters P80; Saint-Gobain Abrasives Pty. Ltd., Thomastown, Victoria, Australia) were placed in the cages. Sandpaper strips were removed four days after blood feeding, wrapped in paper towel and placed in zip-lock bags. The next day, sandpaper strips were labelled and placed in a single sealed chamber with a saturated solution of potassium chloride to maintain the relative humidity ∼80%. The chamber was placed in a controlled temperature room at 26°C with a 12:12 light:dark cycle. To measure quiescent egg viability, batches of >40 eggs were removed from the chamber and submerged in 500 mL rectangular plastic trays filled with 300 mL of water. Each tray was provided with a small amount of fish food (TetraMin tropical fish food tablets, Tetra, Melle, Germany) and a few grains of yeast to stimulate hatching and provide food for larvae. Egg hatch proportions were scored at least three days after hatching by dividing the number of hatched eggs (with a clearly detached cap) by the total number of eggs in each batch. Trays that were not scored for egg hatch after 3 d were stored at 4°C to prevent further hatching and larval development. For all experiments, batches of eggs were hatched on weeks 1 and 2, then every two weeks until week 24. The number of replicate batches of eggs per population and timepoint varied between experiments (see below).

### Variation in quiescent egg viability across laboratory *Aedes aegypti* populations

We performed two experiments to assess variation in quiescent egg viability in *w*Mel and uninfected laboratory and near-field populations. The first experiment was performed in November 2018 with Lab (∼F_60_) and YK (F_11_) populations, both *w*Mel-infected and uninfected. Each population (*w*Mel Lab, uninfected Lab, *w*Mel YK F_11_ and uninfected YK F_11_) was maintained as two separate replicate lines. We scored egg viability for six replicate batches of eggs per time point, per replicate line, for a total of 12 replicate batches per population at each time point. In August 2019, we performed a second experiment with a single replicate line of each population from the first experiment (*w*Mel Lab, uninfected Lab, *w*Mel YK F_18_ and uninfected YK F_18_. We also included additional populations (both *w*Mel-infected and uninfected) collected from Yorkeys Knob and Gordonvale in April 2019 which were at F_3_ at the time of the experiments, and a population from central Cairns at F_11_. In this experiment, eight replicate batches of eggs were tested per population, per time point.

### Effects of laboratory adaptation on quiescent egg viability

To further investigate the effects of laboratory rearing and *w*Mel infection on quiescent egg viability, we performed a third experiment to compare populations collected at two time points from two locations. *w*Mel-infected populations were collected from Yorkeys Knob and Gordonvale in April 2019 and March 2020 and uninfected populations were generated according to the procedure above (see “antibiotic curing”). The experiment was performed in August 2020, with the *w*Mel and uninfected YK and GV populations at F_4_ and F_15_ in the laboratory at the time of experiments. In this experiment, eight replicate batches of eggs were tested per population, per time point. Data for weeks 10 and 20 were discarded due to fungal growth preventing accurate scoring of egg hatch proportions following cold storage.

### *w*Mel origin and genetic background contributions to quiescent egg viability

To estimate the contribution of *w*Mel origin and genetic background to quiescent egg viability, we performed reciprocal backcrossing between *w*Mel Lab and *w*Mel-infected populations from Yorkeys Knob and Gordonvale collected in March 2020. One hundred females from each population (*w*Mel YK, *w*Mel GV and *w*Mel Lab) were crossed to 100 males from each population (*w*Mel YK, *w*Mel GV and *w*Mel Lab). Female progeny were crossed to non-backcrossed males from the same population for two additional generations. This resulted in the partial introgression of *w*Mel from each origin (Lab, Yorkeys Knob and Gordonvale) to the target background (Lab, Yorkeys Knob and Gordonvale), with an estimated 87.5% similarity to the target background. Backcrosses were performed after the first generation of laboratory rearing so that populations were at F_4_ at the time of experiments. This set of comparisons was performed as part of experiment 3, therefore data for *w*Mel Lab, *w*Mel YK F_4_ and wMel GV F_4_ (the non-backcrossed populations) were included in two sets of analyses.

### Cytoplasmic incompatibility and female infertility

We tested cytoplasmic incompatibility and female infertility in *w*Mel-infected individuals hatching from long-term quiescence. Larvae hatching from *w*Mel Lab, *w*Mel YK F_4_ and wMel GV F_4_ eggs stored for 1, 20 or 24 weeks were reared to the pupal stage and separated by sex. We then performed the following crosses: 1. stored *w*Mel-infected females were crossed to non-stored uninfected males, 2. non-stored uninfected females were crossed to stored *w*Mel-infected males, and 3. stored *w*Mel-infected females were crossed to non-stored wMel-infected males. This design allowed us to test for 1. effects of storage on female fertility, 2. effects on storage on the strength of cytoplasmic incompatibility induced by *w*Mel-infected males and 3. effects of storage on the compatibility of *w*Mel-infected females with *w*Mel-infected males. Uninfected females stored for 1 or 20 weeks were also crossed to non-stored uninfected males to test for effects of egg storage on female fertility in the absence of *w*Mel. Females in each cross were blood fed and isolated for oviposition 5 days after crosses were established. Four days after blood feeding, eggs were collected on sandpaper strips from individual females, partially dried, then hatched 3 days after collection. We scored fecundity by counting the total number of eggs laid per female and scored egg hatch proportions according to the procedure above for quiescent egg viability. Females that did not lay eggs were scored as infertile. Up to 20 individuals from each *w*Mel-infected population were measured per cross, but due to low egg hatch following long-term storage we were left with fewer than 20 individuals per cross for some *w*Mel-infected populations.

### Statistical analysis

Most changes in egg hatch proportions were evident from a visual evaluation of the populations across the egg storage period and we used changes across time in comparing across populations. However we were also interested in comparing the size of different effects on hatch rates (including laboratory rearing, *w*Mel infection and genetic background) and for this purpose we focused on two period of storage between 1 and 8 weeks when there was a relatively consistent decline in viability, and between 12 and 18 weeks when population effects were strong based on preliminary experimental data. We also had relatively complete data sets for these periods. We computed logits for all the independent egg hatch proportion measures [33] and then ran general linear models comparing populations across these periods including weeks in storage as a covariate to reflect the continued decrease in hatch rates. We included interaction terms with weeks in storage in the models testing for population effects. General linear models were analysed with IBM SPSS Statistics 26. For the population term, we computed the effect size (partial Eta squared) to allow comparisons between periods and treatments. For the assays on cytoplasmic incompatibility, the data was usually highly skewed towards near-complete hatching or no hatching. We therefore undertook pairwise comparisons between relevant treatments with non-parametric Mann-Whitney U tests. Egg hatch proportions did not differ between different *w*Mel-infected populations within the same cross according to Mann-Whitney U tests, so these were pooled for analysis. For the fertility comparisons we used Fisher’s exact tests using RStudio version 2022.02.2 and the rcompanion package [34] to compare the number of infertile and fertile females across the populations for each time point tested.

## Results

### Variable effects of *w*Mel infection on quiescent egg viability in *Aedes aegypti* laboratory populations

We tested the quiescent egg viability of *Ae. aegypti* populations that were either *w*Mel-infected or uninfected (cured) and had been reared in the laboratory for different numbers of generations. In a pilot experiment (Figure S1), we found that *w*Mel had a weaker cost to quiescent egg viability in a long-term laboratory population (Lab), compared to a population collected from the field more recently (YK F_11_). To further investigate this variability, we collected fresh populations from two locations, Gordonvale (GV) and Yorkeys Knob (YK), which represent areas where *w*Mel was first released in Australia [18]. *w*Mel infection reduced quiescent egg viability in all populations (Figure 1), but there was a high level of variation in the *w*Mel populations compared (GLM, weeks 1-8: F_4,175_ = 30.407, P < 0.001, partial Eta squared = 0.410; weeks 12-18: F_4,140_ = 175.707, P < 0.001, partial Eta squared = 0.834). The main effect of week was also significant in both periods (P < 0.001) and there was a week by population interaction for weeks 12-18 (P < 0.001) but not for weeks 1-8 (P = 0.379). The *w*Mel Lab population had more than twice the egg viability of *w*Mel GV F_3_ at most time points and the other populations fell between these extremes (Figure 1A). In contrast, the five matched uninfected populations showed similar patterns of quiescent egg viability (Figure 1B), with no significant effect of population in weeks 1-8 (F_4,173_ = 0.722, P = 0.578, partial Eta squared = 0.016) but a significant week effect (P < 0.001) but no interaction between week and population (P = 0.975). In weeks 12-18 there was a population effect (F_4,140_ = 7.485, P < 0.001) but this was much smaller (partial Eta squared = 0.176) than for the earlier period and other effects were not significant (P > 0.078). The lack of consistent differences between uninfected populations from the near-field and laboratory counterparts indicate that laboratory adaptation has only a minor effect on this trait. This suggests that the differences between the *w*Mel-infected populations are due to *Wolbachia* infection or its interaction with the genetic background of those populations.

**Figure 1.**
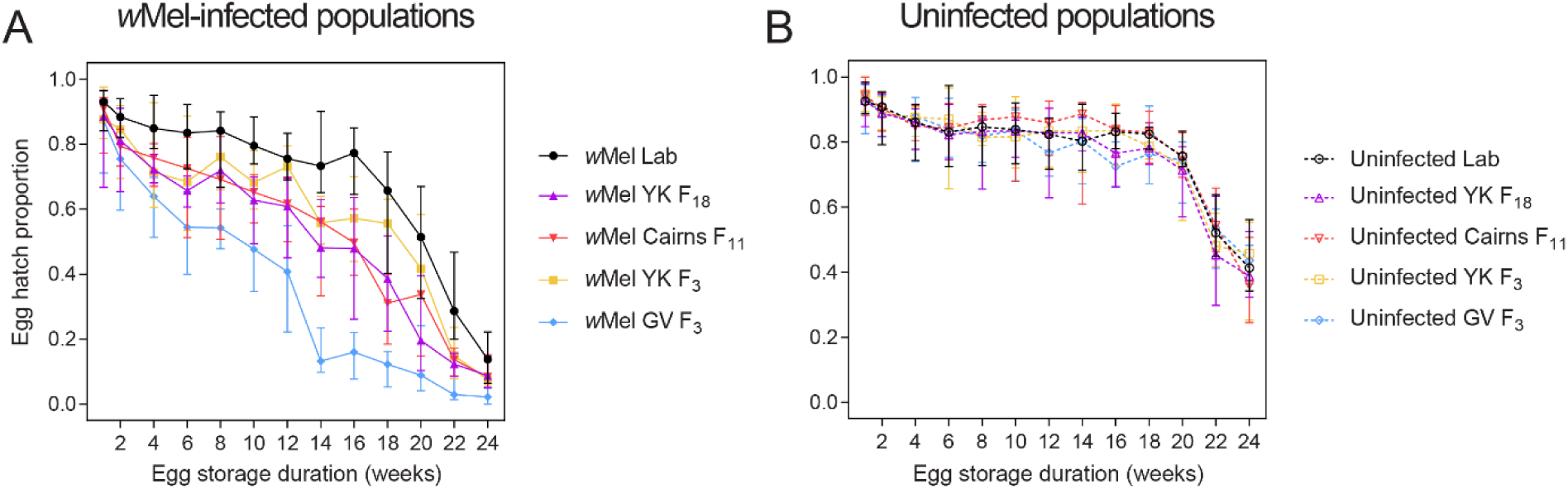
Quiescent egg viability of (A) *w*Mel-infected and (B) uninfected *Ae. aegypti* across different laboratory generations. Symbols and error bars represent median egg hatch proportions and 95% confidence intervals at each duration of egg storage.

### Laboratory adaptation reduces the cost of *w*Mel infection to quiescent egg viability

To further investigate the effect of laboratory rearing, we collected *w*Mel-infected mosquitoes from both Gordonvale and Yorkey’s Knob at two time points (one year apart) and generated uninfected counterparts. Consistent with the first experiment (Figure 1) and the pilot experiment (Figure S1), the uninfected populations had similar quiescent egg viability (Figure 2), though significant population effects were detected across all uninfected populations (GLM, weeks 1-8: F_4,162_ = 8.607, P < 0.001, partial Eta squared = 0.175; weeks 12-18: F_4,140_ = 8.840, P < 0.001, partial Eta squared = 0.202, Figure S2). *w*Mel-infected populations showed larger differences than uninfected populations (weeks 1-8: F_4,174_ = 66.124, P < 0.001, partial Eta squared = 0.603; weeks 12-18: F_4,139_ = 72.650, P < 0.001, partial Eta squared = 0.676). There were no significant interactions between week and population in these comparisons (all P > 0.154). In both the Yorkeys Knob (Figure 2A) and Gordonvale (Figure 2B) populations, quiescent eggs from the F_15_ populations had higher viability than those from the F_4_ populations collected more recently from the field. The effects of *Wolbachia* infection were stronger in F_4_ populations (weeks 1-8: F_1,138_ = 345.048, P < 0.001, partial Eta squared = 0.714; weeks 12-18: F_1,111_ = 773.583, P < 0.001, partial Eta squared = 0.875) compared to F_15_ populations (weeks 1-8: F_1,135_ = 108.100, P < 0.001, partial Eta squared = 0.445; weeks 12-18: F_1,112_ = 345.214, P < 0.001, partial Eta squared = 0.755). This suggests that adaptation to laboratory conditions reduces the cost of the *w*Mel infection to quiescent egg viability.

**Figure 2.**
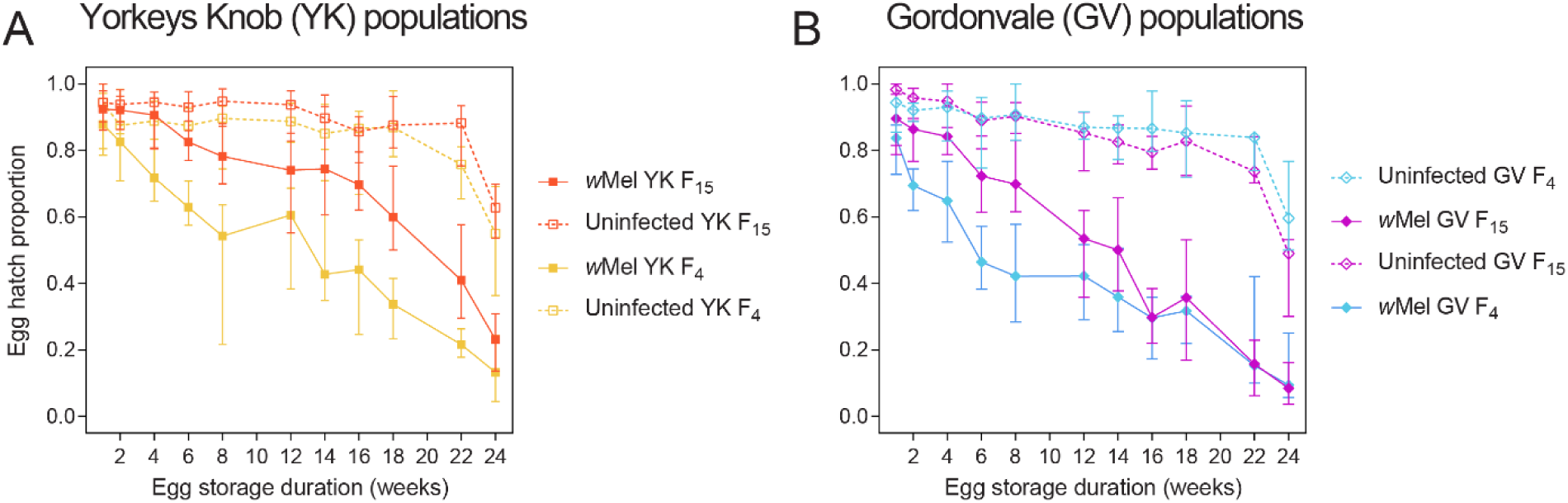
Effects of laboratory rearing on quiescent egg viability in *Ae. aegypti* populations from (A) Yorkeys Knob and (B) Gordonvale. Populations were either *w*Mel-infected (solid lines) or uninfected (dashed lines). Symbols and error bars represent median egg hatch proportions and 95% confidence intervals at each duration of egg storage.

### Costs of *w*Mel infection to quiescent egg viability depend on genetic background and not *Wolbachia* origin

We performed reciprocal backcrosses between the *w*Mel YK, *w*Mel GV and *w*Mel Lab populations to introduce wMel from different origins to different genetic backgrounds and assess the contributions of *Wolbachia* origin and background to quiescent egg viability. When *Wolbachia* from different origins was introduced to the same background, we found similar effects on quiescent egg viability (Figure 3A-C). In contrast, *w*Mel had consistent differences in costs when introduced to different backgrounds (Figure 3D-F). Regardless of the origin of the *w*Mel infection, *w*Mel-infected populations with a lab background performed better than the YK background, which performed better than GV. Overall, the effects of background (weeks 1-8: F_2,311_ = 157.361, P < 0.001, partial Eta squared = 0.503; weeks 12-18: F_2,249_ = 238.517, P < 0.001, partial Eta squared = 0.657) were much larger than those of *w*Mel origin (weeks 1-8: F_2,311_ = 19.234, P < 0.001, partial Eta squared = 0.110; weeks 12-18: F_2,249_ = 27.036, P < 0.001, partial Eta squared = 0.178). Differences between YK and GV suggest adaptation to local conditions that differ between these locales, and a pattern that is consistent with that of the earlier experiments (Figure S1, Figure 2) showing that costs are weaker in lab backgrounds compared to those of recently established populations. Note that differences between the backgrounds were more substantial for the experiments undertaken with *Wolbachia* from the field populations, however it should be noted that replacement of the genetic background was incomplete (estimated at 87.5%) so the *w*Mel origin of the different backgrounds also included a residual nuclear component.

**Figure 3.**
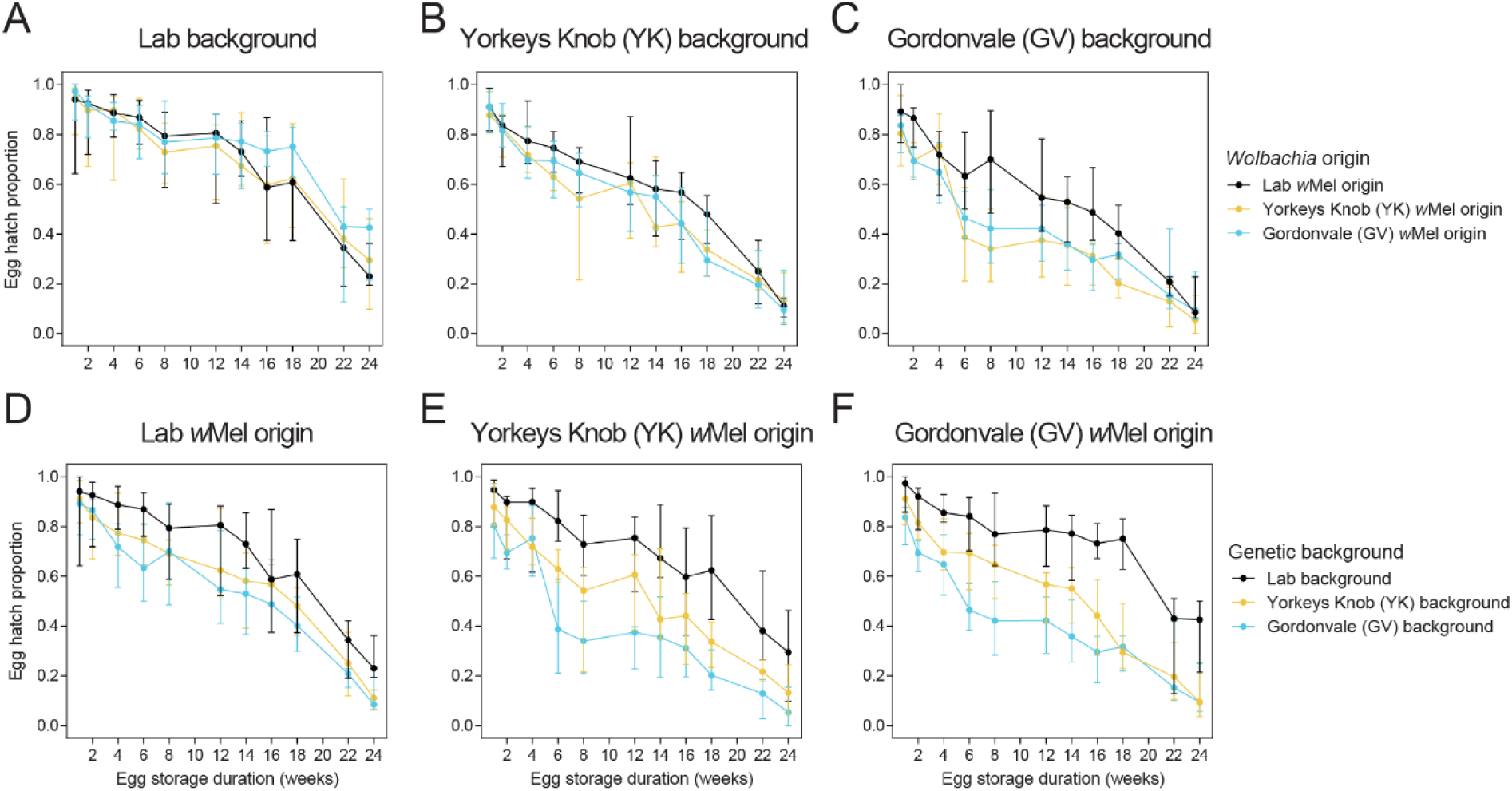
Effects of *Wolbachia* origin (A-C) and genetic background (D-F) on quiescent egg viability in reciprocally-backcrossed *w*Mel-infected *Aedes aegypti*. Panels A-C show comparisons between populations with *w*Mel from Lab (black), Yorkeys Knob (yellow) or Gordonvale (blue) origins when introduced into the same genetic background: Lab (A), Yorkeys Knob (B) or Gordonvale (C). Panels D-F show comparisons between populations with *w*Mel from a single origin: Lab (D), Yorkey’s Knob (E) or Gordonvale (F) that has been introduced to the genetic backgrounds from the Lab (black), Yorkey’s Knob (Yellow) and Gordonvale (blue) populations. Symbols and error bars represent median egg hatch proportions and 95% confidence intervals at each duration of egg storage.

### Long-term egg storage weakens cytoplasmic incompatibility by *w*Mel-infected males and rescue by *w*Mel-infected females

We measured the ability of *w*Mel-infected mosquitoes to induce cytoplasmic incompatibility following long-term egg storage. In non-stored eggs, *w*Mel induced complete cytoplasmic incompatibility (no eggs hatching) when compared to controls that were all compatible, mostly to a high level (Figure 4A). When eggs were stored for 20 (Figure 4B) or 24 (Figure 4C) weeks, the cytoplasmic incompatibility induced by *w*Mel-infected males was incomplete, with uninfected females producing some viable progeny across both time points. Egg hatch proportions in the incompatible cross were significantly higher when males were stored for 20 (Mann-Whitney U: Z = 2.750, P = 0.006) but not 24 (Z = 1.047, P = 0.294) weeks compared to 1 week. *w*Mel-infected females also showed a reduced ability to restore compatibility with *w*Mel-infected males that had not been stored, with reduced hatch proportions when *w*Mel stored females were crossed to *w*Mel males compared to uninfected males (Mann-Whitney U: 20 weeks: Z = 2.750, P = 0.006, 24 weeks: Z = 4.307, P < 0.001).

**Figure 4.**
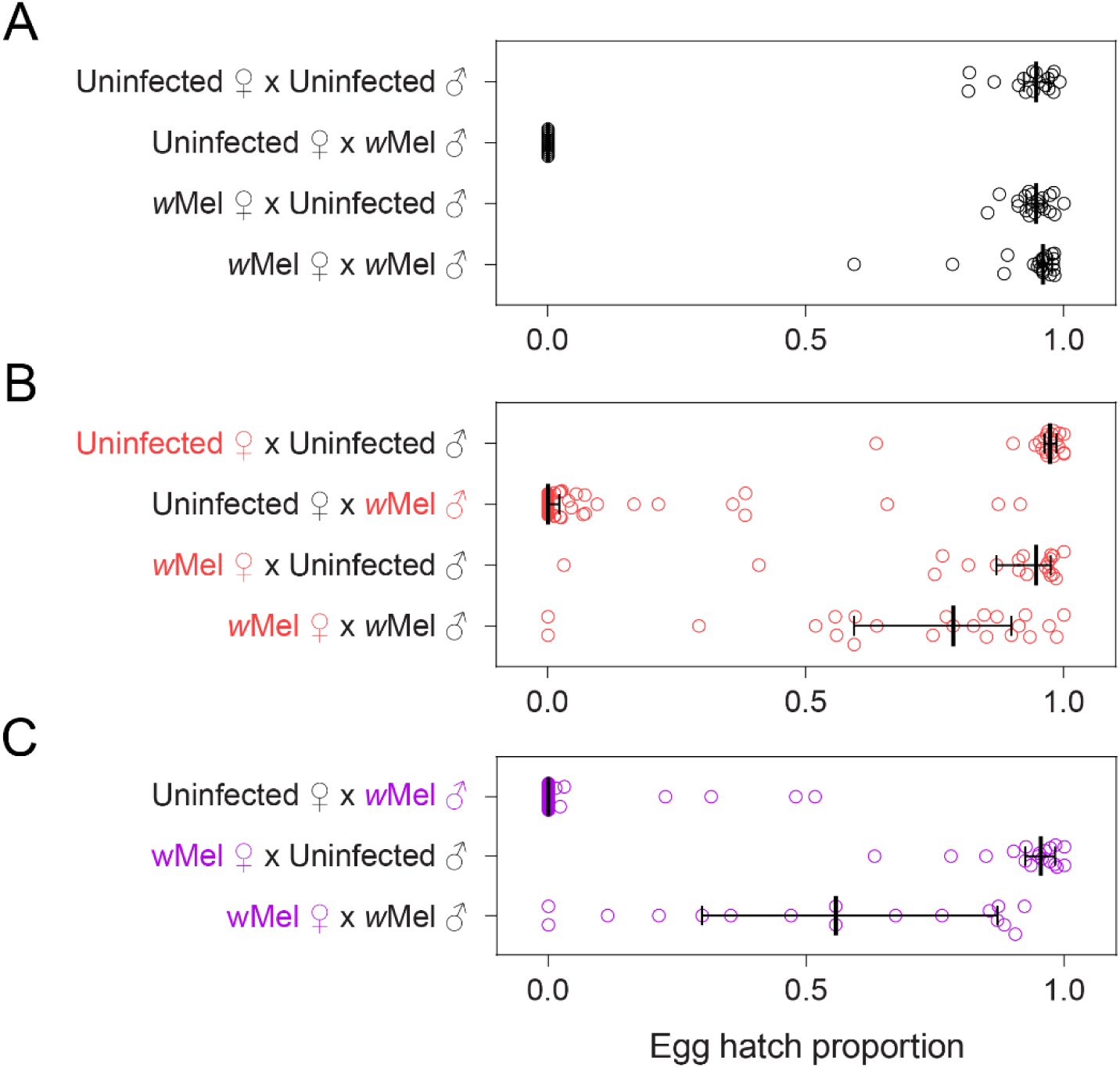
Cytoplasmic incompatibility and compatibility restoration in *w*Mel-infected *Aedes aegypti* hatching from 1 (A), 20 (B) and 24 (C) weeks of egg quiescence. Coloured text indicates sexes that were stored as eggs for 20 (red) or 24 (purple) weeks. Dots show egg hatch proportions for individual females while vertical lines and error bars show medians and 95% confidence intervals.

### *w*Mel-infected females hatching from long-term stored eggs become infertile

We showed previously that the *w*AlbB *Wolbachia* strain causes female mosquitoes to become infertile if they hatch from stored eggs [13, 35]. We scored the proportion of females used in crosses in Figure 4 that laid eggs as a measure of infertility, although we note that the storage period to induce this effect is much longer than described previously for *wA*lbB [13].

The data show a dramatic effect of egg storage on fertility, much stronger than the effects on incompatibility described in the previous section. About half the *w*Mel females stored for 20 or 24 weeks did not lay eggs (Table 1). Across all *w*Mel-infected populations, the proportion of infertile females was 0.54 (binomial confidence interval: 0.44-0.65, n = 94), while for 24 weeks it was 0.51 (0.40-0.63, n = 76). In contrast, no infertile females were observed for any uninfected population or for any *w*Mel population stored for 1 week (upper binomial confidence interval: 0.17).

**Table 1.** Proportion of females from wMel-infected and uninfected *Aedes aegypti* populations hatching from stored eggs that did not lay eggs (i.e. were infertile).

| Egg storage duration | Population | Proportion infertile (n tested)* | Fisher's exact test P value* |
| --- | --- | --- | --- |
| 1 week | Uninfected Lab | 0 (97) a | - |
|  | wMel Lab | 0 (20) a | 1.000 |
|  | wMel YK F<sub>4</sub> | 0 (20) a | 1.000 |
|  | wMel GV F<sub>4</sub> | 0 (20) a | 1.000 |
| 20 weeks | Uninfected Lab | 0 (20) a | - |
|  | wMel Lab | 0.58 (38) bc | < 0.001 |
|  | wMel YK F<sub>4</sub> | 0.39 (36) b | 0.002 |
|  | wMel GV F<sub>4</sub> | 0.75 (20) c | < 0.001 |
| 24 weeks | Uninfected Lab | 0 (20) a | - |
|  | wMel Lab | 0.71 (35) b | < 0.001 |
|  | wMel YK F<sub>4</sub> | 0.30 (33) c | 0.013 |
|  | wMel GV F<sub>4</sub> | 0.50 (8) bc | 0.007 |

\*For each egg storage duration, populations with the same letter are not significantly different by Fisher's exact test, with P values adjusted by the FDR method for multiple comparisons (Benjamini–Hochberg false discovery rate). P values are shown for pairwise comparisons with the Uninfected Lab population.

## Discussion

It is well known that *Wolbachia* infections reduce the viability of quiescent *Ae. aegypti* eggs, including the *w*Mel strain [36]. Here we show that this cost has shifted in mosquito populations that are adapted to different environments. Differences in egg viability were not due to genetic background or *Wolbachia* infection alone; uninfected populations showed similar patterns of egg viability and *Wolbachia* infections from different origins had similar effects when introduced to the same background. However, we found interactions between genetic background and *Wolbachia* infection which likely reflect co-adaptation between *Wolbachia* and mosquito. Our study provides some of the first direct evidence for adaptation of *Aedes aegypti* to *Wolbachia* infection in natural populations.

Post-release monitoring of mosquito populations is an important part of any *Wolbachia* release program [37]. Previous studies have monitored the phenotypic effects of *Wolbachia* in field-collected populations and found them to be largely consistent with pre-release laboratory populations. Gesto et al. [38] and Ahmad et al. [39] show an increase in *Wolbachia* density and sustained virus blocking after *Wolbachia* releases in Brazil (*w*Mel) and Malaysia (*w*AlbB) respectively. In Cairns, Australia, post-release monitoring found persistent costs of the *w*Mel infection to female fertility [26] and sustained virus blocking [40] after one year. Phenotypic effects have largely persisted in the longer term, with one exception being development time where costs were only apparent in a laboratory background [23]. However, no studies to date have identified the basis of population differences. Here we used a combination of repeated field collections, antibiotic curing and reciprocal backcrossing to show that fitness costs depend on genetic background and shift with laboratory adaptation. This finding is consistent with previous studies demonstrating the potential for host genetic changes to mediate fitness costs [15, 41] and the lack of changes in the *w*Mel genome itself [21-23]. We have previously found that the genomes of *Ae. aegypti* have not changed much across the 10 year period since releases of *w*Mel started, although some outliers were detected that may reflect adaptive responses [24]. The current study is the first case where shifts in fitness costs have been linked to host genetic changes under field conditions.

The patterns of quiescent egg viability costs we observed were puzzling, given that we found higher costs of *w*Mel in field-collected populations which experience long dry seasons (www.bom.gov.au) and where selection for increased quiescent egg viability is expected and the costs of *w*Mel should attenuate. Costs were unexpectedly weaker in laboratory-adapted populations where eggs were usually stored for less than two weeks, therefore with little selection to maintain long-term quiescent egg viability. It is possible that these patterns are due to trade-offs, where *w*Mel provides a benefit under field conditions at the cost of reducing quiescent egg viability. In natural *Wolbachia*-insect associations, *Wolbachia* infections often provide context-dependent fitness costs and benefits [20].

*Wolbachia* infections typically induce greater fitness costs in novel mosquito hosts than in natural hosts, but these costs are expected to attenuate over time [42]. In a previous study, we found that the costs of *w*Mel to quiescent egg viability were consistently low when *w*Mel was introduced to a naïve laboratory genetic background [23]. While the experiments are not directly comparable, together they suggest that costs may depend more on local mosquito adaptation than the novelty of the *Wolbachia*-mosquito association. It is therefore possible that our findings reflect innate differences in mosquito populations rather than adaptation to *Wolbachia* infection, but confirming this would require *Wolbachia*-free field populations which are no longer present in our study region due to widespread *w*Mel coverage [7, 43]. Future work investigating adaptation of mosquitoes to *Wolbachia* infection should consider effects in both naïve and “*Wolbachia*-adapted” backgrounds and in both field- and laboratory-adapted populations.

Our study identified novel and substantial fitness costs of *w*Mel infection following long-term storage of eggs. When eggs from *w*Mel-infected populations were stored for extended periods (20 or 24 weeks), males partially lost their ability to induce cytoplasmic incompatibility and around half of all females became infertile, with fertile females partially losing their compatibility with *Wolbachia* infected males. This effect was *Wolbachia*-specific because uninfected females stored for the same amount of time never became infertile. The effects of egg storage on infertility are similar to those described previously for the *w*AlbB infection [13, 35] but substantially weaker, with ∼50% infertility observed after only nine weeks for *w*AlbB. While the additional costs of *w*Mel infection were only apparent after long-term storage, interactions with other environmental factors such as high temperatures, which also reduce cytoplasmic incompatibility by *w*Mel [44], could exacerbate these costs. These cumulative effects on fitness may influence *Wolbachia* spread and persistence in locations with long dry seasons where these costs will be most apparent.

Host adaptation to novel *Wolbachia* infection is poorly understood and our study provides a foundation for future investigations. It is worth exploring host adaptation to *Wolbachia* strains with more substantial effects on fitness, for instance *w*AlbB which is established in natural populations in Malaysia [8]. The consistent differences in *w*Mel effects between Yorkey’s Knob and Gordonvale populations also highlight the potential for local adaptation to influence fitness costs; a factor which is rarely considered [45]. Host genotype effects on fitness costs have important implications for *Wolbachia* releases given that the strength of fitness costs drives *Wolbachia* establishment potential [14]. Our work also prompts further investigation into the extent to which adaptation to artificial rearing conditions masks the fitness costs of *Wolbachia*, particularly under stressful conditions where *Wolbachia* infections have clear costs and where selective pressures differ greatly between the laboratory and the field. Our results here suggest that fitness comparisons between mosquitoes with laboratory genetic backgrounds may not always predict *Wolbachia* fitness costs in wild mosquito populations. When planning a local *Wolbachia* release program it is therefore important to use locally-sourced mosquitoes that have spent minimal time in the laboratory for phenotypic assessments.

## Acknowledgements

We thank Marianne Coquilleau for assistance with experiments and Ashley Callahan and Qiong Yang for routine screening of *Wolbachia*-infected mosquito populations. We also thank Kyran Staunton and Scott Ritchie from James Cook University, Cairns for providing field-collected *Aedes aegypti*.

## Supplementary information

**Figure S1.**
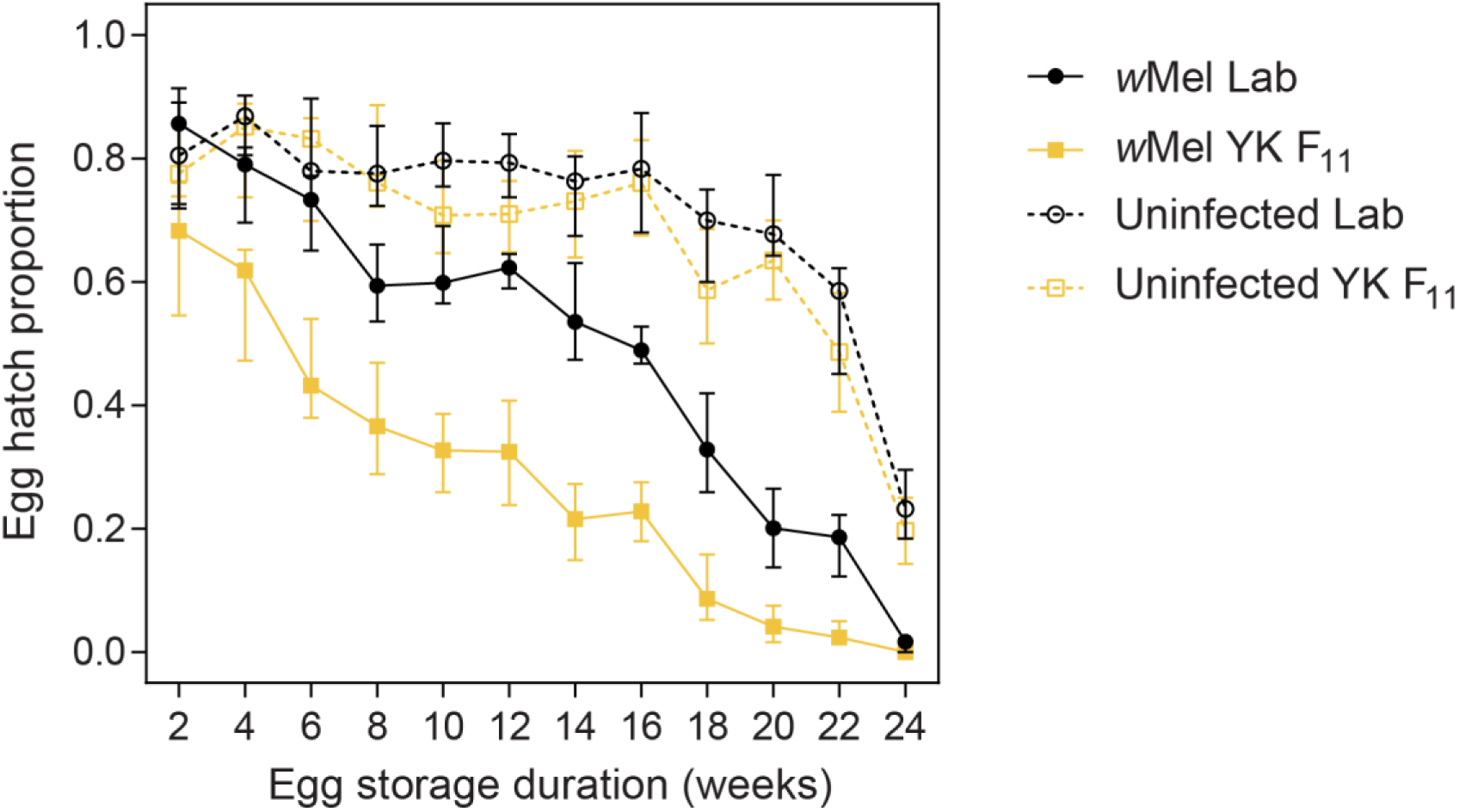
Quiescent egg viability of *w*Mel-infected and uninfected *Ae. aegypti* from laboratory (Lab) and Yorkeys Knob (YK F_11_) populations. Data are pooled from two replicate populations. Symbols and error bars represent median egg hatch proportions and 95% confidence intervals at each duration of egg storage.

**Figure S2.**
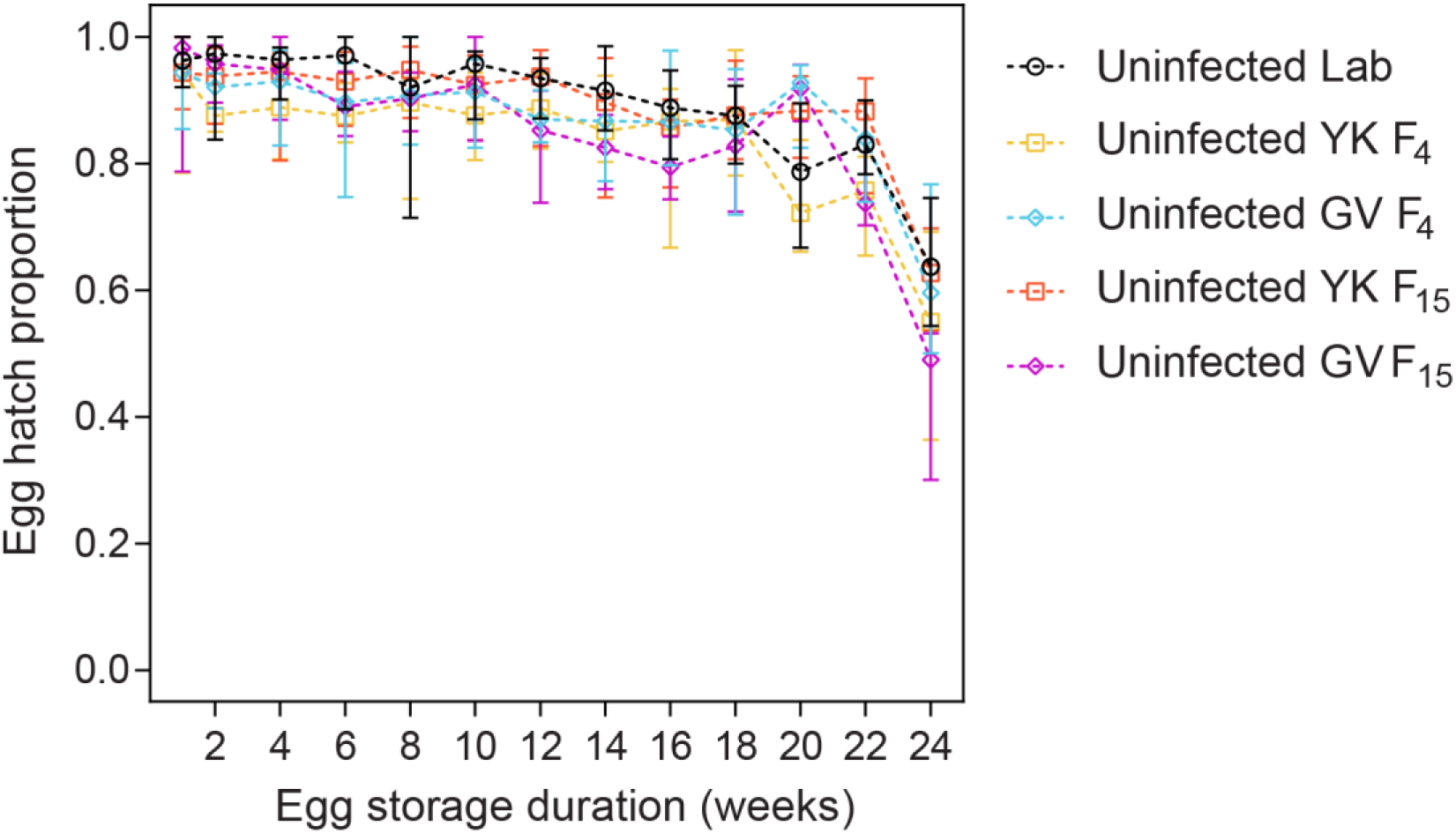
Quiescent egg viability of uninfected *Ae. aegypti* populations included in experiments to test effects of laboratory rearing and genetic background. Symbols and error bars represent median egg hatch proportions and 95% confidence intervals at each duration of egg storage.

## Notes

### Competing Interest Statement

The authors have declared no competing interest.

